# Multilevel Modelling of Gaze from Hearing-impaired Listeners following a Realistic Conversation

**DOI:** 10.1101/2022.11.08.515622

**Authors:** Martha M. Shiell, Jeppe Høy-Christensen, Martin A. Skoglund, Gitte Keidser, Johannes Zaar, Sergi Rotger-Griful

**Affiliations:** Eriksholm Research Centre, Oticon A/S, Snekkersten, Denmark; Division of Automatic Control, Department of Electrical Engineering, The Institute of Technology, Linköping University, Linköping, Sweden; Department of Behavioural Sciences and Learning, Linneaus Center HEAD, Linköping University, Linköping, Sweden; Hearing Systems Section, Department of Health Technology, Technical University of Denmark, Kgs. Lyngby, Denmark

## Abstract

**Purpose:** There is a need for outcome measures that predict real-world communication abilities in hearing-impaired people. We outline a potential method for this and use it to answer the question of when, and how much, hearing-impaired listeners look towards a new talker in a conversation.

**Method:** Twenty-two older hearing-impaired adults followed a pre-recorded two-person audiovisual conversation in the presence of babble noise. We compared their eye-gaze direction to the conversation in two multilevel logistic regression (MLR) analyses. First, we split the conversation into events classified by the number of active talkers within a turn or a transition, and we tested if these predicted the listener’s gaze. Second, we mapped the odds that a listener gazed towards a new talker over time during a conversation transition.

**Results:** We found no evidence that our conversation events predicted changes in the listener’s gaze, but the listener’s gaze towards the new talker during a silent-transition was predicted by time: The odds of looking at the new talker increased in an s-shaped curve from at least 0.4 seconds before to 1 second after the onset of the new talker’s speech. A comparison of models with different random effects indicated that more variance was explained by differences between individual conversation events than by differences between individual listeners.

**Conclusion:** MLR modelling of eye-gaze during talker transitions is a promising approach to study a listener’s perception of realistic conversation. Our experience provides insight to guide future research with this method.

## Introduction

Difficulty with everyday conversation is a debilitating consequence of hearing loss that is not well captured by traditional audiometric tests (Kiessling et al., 2003). In recent years, the field of hearing science has increasingly emphasized the need for improved ecological validity in research outcomes for a better understanding of the real-world challenges of hearing loss (Keidser et al., 2020). Numerous new research paradigms have been implemented that aim to tap into more complex behaviors and environments (for a review, see Carlile & Keidser, 2020). In the current investigation, we continue this agenda by studying the behavior of hearing-impaired listeners following a realistic, two-person, audiovisual conversation. One challenge of working with this type of stimuli is that the realism of the stimulus comes at a cost to experimental control. As such, reliable and sensitive measures of the listener’s experience with conversation stimuli are needed. In the current investigation, we introduce a novel approach for this: Multilevel logistic regression modelling of the listener’s eye-gaze during a conversational transition between the talkers. Our purpose is to outline this paradigm as a potential tool for understanding the behavior of hearing-impaired listeners in realistic conversation, so as to inform the growing research interest in this topic.

In the study of language comprehension, the measurement of where someone looks has been widely used as a proxy for underlying cognitive processes via the so-called visual world paradigm (Huettig et al., 2011). The approach evolved from the observation that listeners move their eyes in characteristic patterns in response to speech stimuli that are accompanied by visual images (Cooper, 1974). With the increased accessibility of eye-tracking technology in the late twentieth century, the use of eye-tracking to understand cognition has bloomed across the social sciences. In the past fifteen years, eye-gaze has also become a topic of interest in the field of hearing science due to its potential application as a signal for controlling hearing aid technology (e.g. Best et al., 2017; Favre-Félix et al., 2018; Hart et al., 2009; Hládek et al., 2019; Kidd, 2017; Roverud et al., 2017; Skoglund et al., 2022). Most recently, researchers have started studying eye-gaze to describe the behavior of hearing-impaired people in realistic multitalker situations (Hadley et al., 2019; Hendrikse & Llorach, 2019; Lu et al., 2021), an application that capitalizes on our innate tendency to look at the targets of our attention and our attraction towards social stimuli.

Faces have long been known to attract visual attention in realistic scenes (e.g. Buswell, 1935), see Hessels, 2020for a review). Compared to when they are static, faces attract even more gaze when they dynamically produce speech (Foulsham & Sanderson, 2013; Scott et al., 2019; Vo et al., 2012). When additional talkers are added to a scene, the reported amount of gaze towards the talkers remains high (Dawson & Foulsham, 2022; Foulsham et al., 2010; Foulsham & Sanderson, 2013; Hidalgo et al., 2022; Hirvenkari et al., 2013), but the duration of the fixations towards them decreases (Foulsham et al., 2010; Šabić et al., 2020). Of interest in these multitalker scenarios though is how the listeners divide their gaze between the talkers over time. In natural conversation, talkers take turns speaking, such that the majority of the time there is only one active talker (Sacks et al., 1974). Unsurprisingly, when viewing multitalker situations, listeners with normal hearing spend the majority of their time gazing at the active talker (Casillas & Frank, 2012; Dawson & Foulsham, 2022; Foulsham et al., 2010; Foulsham & Sanderson, 2013; Hidalgo et al., 2022; Hirvenkari et al., 2013; Tice & Henetz, 2011; Vertegaal et al., 2001).

Gazing at the active talker in a conversation requires that a listener executes a high-speed eyemovement – a saccade - to direct their gaze from one talker to the next. Saccades in response to speech stimuli require 200 ms to plan (Matin et al., 1993). Since a typical gap between talker’s turns in a conversation also lasts 200 ms (Stivers et al., 2009), the listener must accurately recognize the end of the previous talkers’ turn if their gaze is to arrive on the new talker in time. Although turn-ends can be deduced from unimodal visual cues alone (Latif et al., 2017), a precise prediction of the timing of a turn-end may be dependent on cues uniquely from the auditory modality (Latif et al., 2018), such as information on syntax, pragmatics, and prosody. As such, if audio degradation from hearing loss reduces turn-end perception, it may also be expected to impair a listener’s ability to precisely follow changing talker activity over time. For this reason, we expect that the eye-gaze of listeners during talker transitions may give insight into the listener’s ability to follow the conversation.

Eye-gaze has been used previously to study the perception of turn-end cues in normal-hearing adults and children (Augusti et al., 2010; Casillas & Frank, 2012; Dawson & Foulsham, 2022; Foulsham & Sanderson, 2013; Hidalgo et al., 2022; Hirvenkari et al., 2013; Holler & Kendrick, 2015; Keitel et al., 2013; Tice & Henetz, 2011; von Hofsten et al., 2009). Generally, the method entails tracking a listener’s eye movements while they view a talker transition in a conversation stimulus, and calculating how the timing of the listener’s saccade from one talker to the next aligns with the corresponding audio signals. This research demonstrates that children as young as three years old anticipate a talker transition by moving their eyes to the new talker (Keitel et al., 2013), and that this behavior can occur in normal-hearing adults even when the conversation is presented unimodally visually (Dawson & Foulsham, 2022; Foulsham & Sanderson, 2013; Hirvenkari et al., 2013), or in an unknown language (Casillas & Frank, 2012). This robustness makes this behavior an ideal target for working with realistic stimuli: In theory, it will occur despite the many different factors that influence realistic conversation.

Promisingly, there is also evidence that the way that the listener’s gaze unfolds over time during a talker transition is sensitive to differences in the listener’s perception. For example, listeners with normal hearing show more early saccades to the new talker when the transition involves a question-and-answer exchange than when it does not (Casillas & Frank, 2012), and saccades are delayed and reduced when the audio of the conversation is muted (Dawson & Foulsham, 2022; Foulsham et al., 2010; Hirvenkari et al., 2013; Tice & Henetz, 2011). Furthermore, in listeners with normal hearing, the first saccade to a new talker in a conversation transition is delayed when the audio is degraded by vocoding, and in some conditions, this saccade’s latency predicts the listener’s comprehension (Hidalgo et al., 2022).

The current investigation builds on the work outlined above by using eye-gaze behavior to study turn-end perception in a new population: hearing-impaired listeners. We introduce a multilevel logistic regression (MLR) modelling approach (Barr, 2008) to this topic, which allows us to provide a formal account of eye-gaze over time during transitions between talkers in a conversation. For an in-depth discussion of the advantages of MLR for eye-gaze analysis, we refer the reader to Barr (2008). In short, we chose this approach because (1) it is sensitive to early changes in gaze behavior (so-called ‘anticipatory effects’, (Barr, 2008), enabling us to capture potentially subtle indications of turn-end perception, (2) its use of hierarchichal groupings of data helps to address the increased variability that can arise between participants and stimuli when targeting ecologically-valid research outcomes, and (3) it can be easily adapted for future research to incorporate comparisons between new conditions, populations, and individual differences. With this MLR approach, we address the research question of when, and how much, hearing-impaired listeners look towards a new talker. Furthermore, we examine how much participants and stimuli differ from one another in our design. Through this work we aim to lay a foundation that can advise future research on how the gaze behavior of hearing-impaired listeners reflects their experience of everyday conversation.

## Methods

### Participants

The data were collected as part of a separate experiment that investigated how to use in-ear electrodes to measure eye-movements (Skoglund et al., 2022). The full experiment, including the use of the data here, was approved by the Ethics Committee of the Capital Region in Denmark (H-20030989) and all participants gave informed written consent. Participants were recruited from the Eriksholm Research Centre test participant pool, where they received audiological services and hearing aids in return for their membership. They were financially compensated for their travel to the test site at Eriksholm Research Centre.

Twenty-six adults completed the study, but four were excluded *post hoc* due to insufficient data because of a poor eye-tracking. The final sample consisted of 22 participants (average years old: 69, range years old: 55-84; 9 female and 13 male). Participants had mild to moderate symmetric hearing loss with a mean pure tone average (in dB HL) of 30.5 in the left ear (range: 12-45.8) and 30.1 in the right ear (range: 12-47), with less than 10 difference between the ears for at least 3 out of 4 frequencies, and no more than 20 difference at any frequency (audiograms shown in Figure 1). They were regular users of hearing aids. All were native Danish speakers with no prior experience with the testing materials, and reported that they could see the stimuli clearly at 1.5 metres distance without the use of glasses or contact lenses (Skoglund et al., 2022).

**Figure 1.**
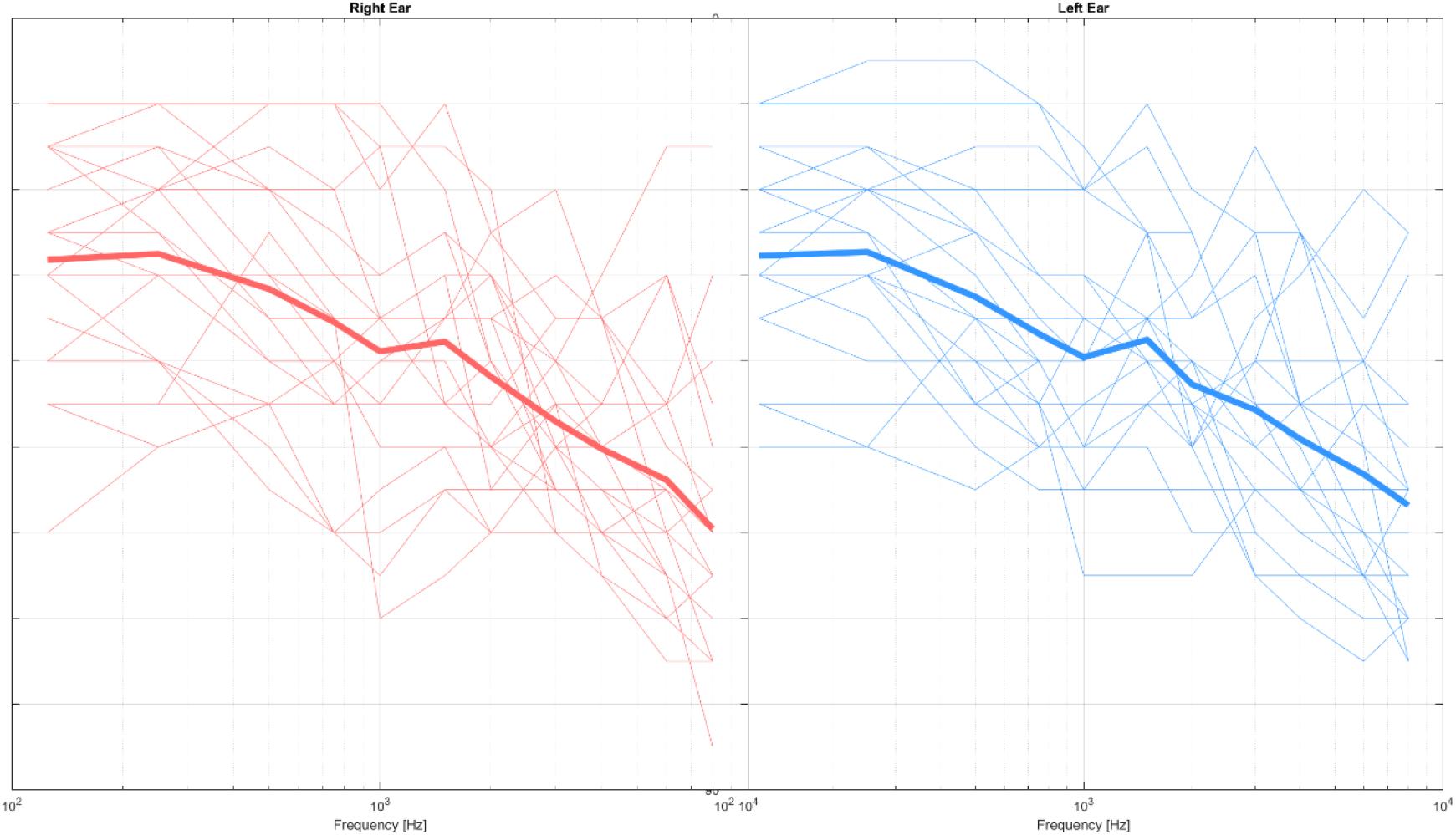
Audiograms for individual participants (thin lines) and group average (thick line).

### Stimuli

Stimuli consisted of audiovisual recordings created at Eriksholm Research Centre. The recordings involved two talkers engaged in an unscripted conversation in Danish. Each talker’s speech was recorded by an individual headworn microphone. We used the Diapix task (Baker & Hazan, 2011) – spot-the-difference between two pictures – to elicit natural turn-taking behavior but still constrain the topic and vocabulary of the conversation. The talkers were seated in chairs facing the video recorder but were free to move their body and head naturally. In this position, they tended to hold their heads slightly angled towards one another, such that only the furthest edge of their lips was occasionally hidden from the viewer. The recordings were cut into clips of conversation ranging in duration of 9-39 s. Every clip finished directly after the talkers uncovered a difference between their pictures.

### Behavioral Task

To motivate the listeners to pay attention to the conversation, a behavioral task was introduced. At the end of each clip, participants were presented with a multi-choice question that asked the listener to identify the type of difference disclosed at the end of the clip. Listeners were given three response options, with the responses consisting of words that did not appear in the talkers’ speech. This was to encourage the listener to attend to the meaning of the speech rather than rely on word recognition strategies.

### Test setup

Videos were shown on an 88” curved Samsung television screen, with audio presented from three loudspeakers (Genelec 8040C), visible on the floor in front of the screen, and seven (Genelec 8010A) mounted in a row above the screen and hidden behind an acoustically-transparent black fabric. The visual stimuli were presented such that the two talkers – shown from head to knee – appeared approximately life-sized on the TV screen, with their heads positioned approximately 20 degrees apart at a viewing distance of 1.5 metres. The audio from each talker was presented from the spatially-corresponding loudspeaker at a sound-pressure level (SPL) of 60 dB. The remaining loudspeakers played a multitalker babble noise, created in-house from a mixture of five male and five female Danish-speaking voices, different from those used for the conversation stimuli, and presented at cumulative level of 60 dB SPL across the loudspeakers. The audio for each loudspeaker was equalized with a finite-impulse response filter based on the inverse of the loudspeakers’ 3^rd^-octave smoothed frequency response function, measured at the center-of-head position. Furthermore, the signal loudness was modified for each participant individually to compensate for their hearing loss in their better ear (Moore & Glasberg, 1998; Skoglund et al., 2022).

A pair of Tobii Pro 2 Eye Tracking Glasses (Tobiipro, Sweden, *https://www.tobiipro.com*) were used to collect eye-movement data that were stored directly on a SD card within the device. The glasses were fitted with a set of six reflective markers that were used to track the position of the eye-tracker in the room via a four-camera motion capture system (Vicon Vero, UK, *https://www.vicon.com*).

Data from the motion capture system was recorded on a desktop computer that also controlled the experiment presentation. Audio was interfaced through two linked soundcards (RME Fireface 400, Hammerfall DSP Multiface II). In addition to connecting to the loudspeakers, the soundcards transmitted a trigger signal to the eye-tracker recording system (via a custom-made trigger box) and to the motion capture system for the purpose of synchronizing the measurements with the audio. Behavioral responses from the participants were recorded through a bluetooth-connected number pad.

The stimuli were controlled with Matlab 2019a (Mathworks, USA, *https://mathworks.com*), which also recorded the participants’ behavioral responses. Matlab interfaced with VLC media player to present the video stimulus, and used msound toolbox (*https://github.com/TGM-Oldenburg/Msound*) to connect to the external soundcard with ASIO drivers. Nexus software (Vicon, UK, *https://vicon.com*) was used to calibrate the motion capture system and record the position of the head. Tobii data were converted to text files with the open-source python toolkit TobiiPyGlassesSuite (De Tommaso & Wykowska, 2019, *https://github.com/ddetommaso/TobiiGlassesPySuite*). All data were processed and analysed with custom scripts in Matlab 2019a and R (version 3.6.1, *https://www.r-project.org*).

### Test Procedure

Each participant was tested in a single visit. The participants were seated in a comfortable chair, positioned such that their head was approximately 1.5 metres from the television screen. After fitting the participant with the eye-tracker and sensor equipment, participants were verbally instructed on the task. The experimenter briefly described the contents of the stimuli and played an example with no background noise. Then the instructions for the trial were given. Each trial began with a fixation cross appearing on a black background, accompanied by the onset of the multitalker babble noise. After three seconds, the conversation speech and video began. The listeners were asked to hold their heads stationary, to look at the fixation cross when it appeared on the screen, to listen to the conversation, and to answer (by button-press) the multiple-choice question that appeared on the screen at the conclusion of each clip. The question and the response options were also explained. The next trial started automatically after the participant’s response. The participants were given 2 blocks of practice trials (19 trials total) and 2 blocks of experiment trials (26 trials total), with the option of breaks in between blocks. These blocks were interleaved with testing blocks for other experimental conditions that were not of interest to the current investigation and are discussed elsewhere (Skoglund et al., 2022).

### Conversation Data Processing

From the individual audio tracks of each talker in our stimuli, we generated voice activity timeseries by segmenting the audio into 5 ms-long windows that overlapped by 1 ms, and using an individually selected root mean square threshold on the segments to remove cross-talk during silent transitions, similar to (Sørensen et al., 2021). In several tracks with substantial cross-talk, periods of the talkers’ silence were manually edited to exclude cross-talk. Individual tracks were normalized to the same energy level for the voice-active portions within a clip, to compensate for variation in the speaking level of the talkers. These voice activity timeseries were used to classify each clips’ timecourse into five different conversation event types. We applied the algorithm from (Sørensen et al., 2021) to identify periods of turns and transitions, and label periods of silence and overlap. From this, the conversation was divided into discrete events based on the number of talkers (either silence, solo, or overlap for 0, 1, or 2 simultaneous talkers) for both turns and transitions separately. This resulted in the following labels, which are illustrated in Figure 2: Silence-turn, silence-transition, solo, overlap-turn, and overlap-transition. Similar to (Heldner & Edlund, 2010), periods of silence-turn that were less than 180 ms in duration were bridged, and voice activity less than 70 ms in duration were excluded. The frequency and durations of the resulting conversation event types can be seen in Table 1.

**Table 1.** Description of conversation stimuli by number of talkers (silence (0), solo (1), or overlap (2)) and turn-taking stage (turn or transition)

| Event Type | No. Occurrences/<br>Participant | Event Duration (s) |  |  |  |
| --- | --- | --- | --- | --- | --- |
|  |  | mean | min. | med. | max. |
| Silence-Turn | 51 | 0.50 | 0.18 | 0.38 | 2.18 |
| Solo | 240 | 0.95 | 0.02 | 0.6 | 7.62 |
| Overlap-Turn | 24 | 0.36 | 0.08 | 0.29 | 1.56 |
| Silence-Transition | 120 | 0.406 | 0.02 | 0.34 | 3.3 |
| Overlap-Transition | 31 | 0.27 | 0.02 | 0.22 | 1 |

**Figure 2.**
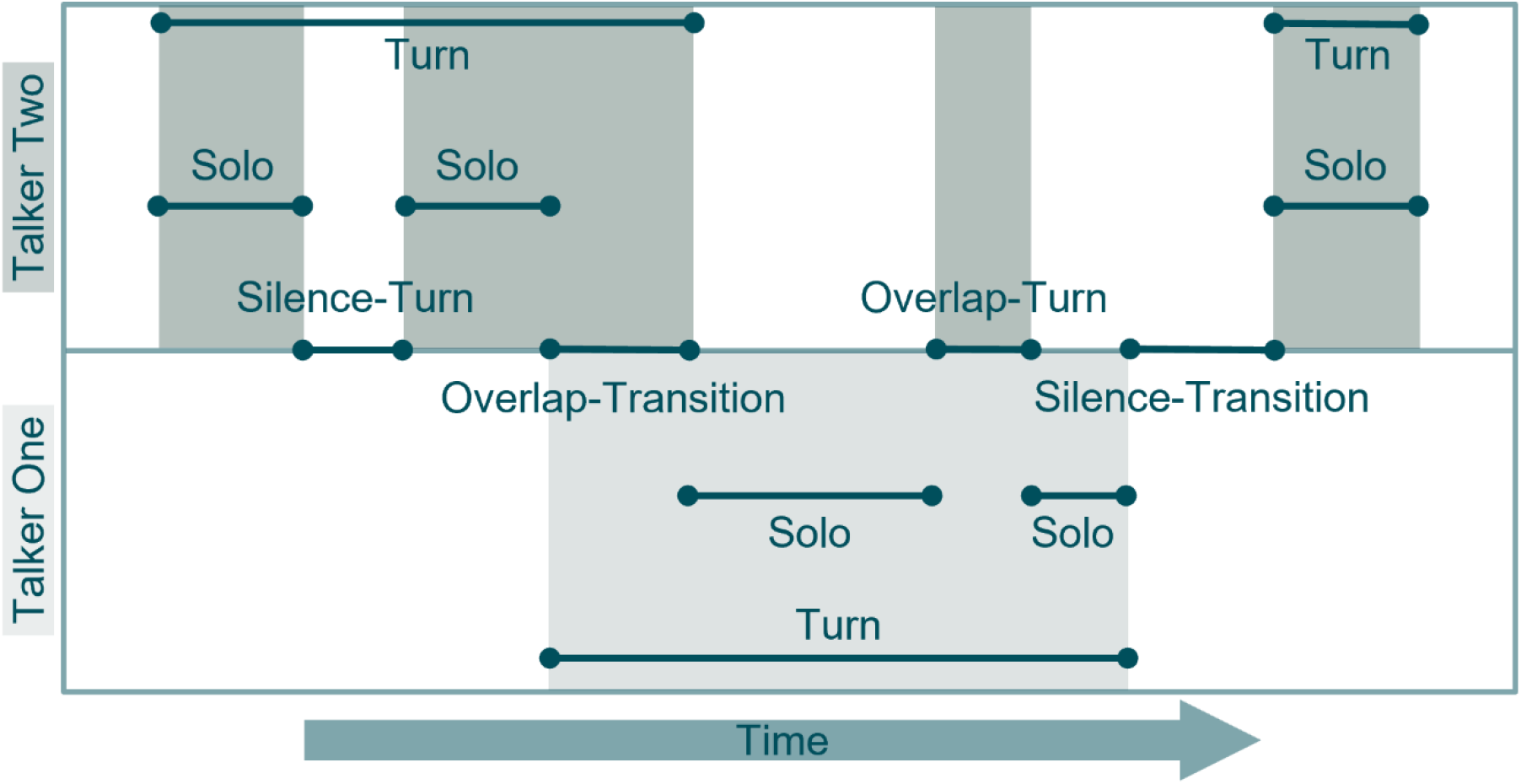
Diagram of conversation event classifications based on the number of active talkers (0, 1, or 2 for silence, solo, and overlap respectively) and turn-taking stage (turn or transition). Shaded boxes indicate voice activity.

To answer our research question, we needed to identify points in the stimuli where we expected listeners to transfer their gaze to the new talker due to a transition in the conversation. Given the low occurrence of overlap-transitions in our stimuli, we selected time periods surrounding the onset of a new talker after silent-transition events only. To exclude transition events that were uncharacteristically long or short, we selected only those in which the durations of the silent period and the subsequent turn were roughly within the interquartile range for each event type (minimum duration of turn for inclusion: 0.4 s, silence: 0.14 s; maximum time included from turn: 2.92 s, silence: 0.508 s). This resulted in a selection of 68 out of the total 120 silence-transition events within our stimuli.

### Eye-gaze Data Processing

Since previous research has established that listeners prefer looking at faces compared to other elements of a scene (see e.g. (Hessels, 2020for a review), our eye-gaze data processing worked from the assumption that our participants did so, and focussed on describing how gaze was divided between talkers in relation to their speech activity. This assumption allowed us to process the data with few *a priori* assumptions, and without physically restricting the participants heads. Our motivation here was to develop a method that can translate well into future experimental setups that have less control over data quality and more stimulus variability, which is a necessary compromise to move into research with even more realistic scenarios. Note that, although we did not compensate for the listener’s head position in our eye-gaze analysis, motion capture data confirmed that participants held their heads mostly stationary during the experiment. Only 4% of trials showed a change in yaw larger than 2 degrees from the average head position in the trial. Only 5% showed translations larger than 14.6 mm on the horizontal azimuth from the average head position in the trial, which corresponds to less than 1 degree of visual angle in our setup. We conclude therefore that head movements made a negligible impact on our analysis.

For each block of trials, we selected the eye that had the lowest proportion of missing data in each participant, and we used the gaze direction coordinates to calculate gaze angle in degrees on the azimuth. We assigned every gaze sample as belonging to one of the two talkers. This assignment was accomplished in an automatic data-driven manner by *k*-means clustering (*k* = 2) applied to the azimuth measurement within each trial. This method ignores that some of the gaze samples belonged to saccades between the talkers, and towards elements of the scene outside of the talkers. We treated these latter gaze samples as inherent noise, and assumed that this noise was not systematically different between the talkers across the full experiment.

Prior to the *k*-means clustering, we removed gaze samples outside of a 50 degree range on the azimuth, centered at the trial’s mean gaze angle. These were assumed to be noise due to their spatial extremeness (keeping in mind that the talker’s were separated by approximately 20 degrees). This cleaning mitigated cluster biases from extreme values. After k-means labelling, we further processed the gaze to conform to physiologically-based assumptions. We assumed that (1) a change in region-of-interest – the moment the gaze crossed the midline between the two talkers - would cover 10 degrees, given that the talkers were approximately 20 degrees apart, that (2) a saccade of this size would last approximately 40 ms, and that (3) a fixation to a talker would last at least 100 ms. Therefore, changes of labels that lasted less than 180 ms were considered noise and were filled in with the previous label. Likewise, chunks of missing data less than 180 ms that did not span a label change were filled in with the previous label.

After this preprocessing, data from 12 out of 572 trials were excluded due to missing more than 40% of the trials’ data. The remaining trials had a total of 8.4% data missing, 3% of which was due to the preprocessing steps described above and the remainder of which was due to signal loss during data collection.

### Target talker labelling

Our modelling analysis required that we first decide which talker to label as the target in each type of conversation event. To do so, we aligned data from all participants by their timestamps, and classified these according to one of the five conversation event types shown in Figure 2 (silence-turn, silence-transition, solo, overlap-turn, overlap-transition). Similar to Hirvenkari et al. (2013) we used a binomial test at each timepoint to test whether the distribution of the group’s gaze was different from random. We identified 10115/15965 timepoints where participants gazed at the same talker (*p* < 0.05, or at least 17/22 participants). We labelled these timepoints according to which of the two targets was selected by the group: finishing vs. starting talker for overlap- and silent-transitions, and turn-holder vs. non-turn-holder for solo, silence-turn, and overlap-turn events. The resulting proportions revealed that the group preferred to look at the finishing talker during transitions and the turn-holder during turns. Specifically, during overlap-transition and silence-transition events, 34.1% and 36.3% of collected timepoints showed that the group looked towards the finishing talker, versus 12.4% and 11.8% towards the starting talker. During solo, overlap-turn, and silence-turn events, 65.1%, 42.8%, and 45.7% of collected timepoints showed that the group looked towards the turn-holder, versus 4.4%, 4.7%, and 0.0% towards non-turn-holder. Based on this, the finishing and turn-holding talkers are herein referred to as the target for any given timepoint. This denotation was used to code the data as either on or off target for the subsequent modelling.

## Results

We tested two different models (Equation 1 & Equation 2) to address our research question. In both, the dependent variable was the probability of gaze-to-target transformed to log odds. We modelled an intercept for each individual transition occurrence (γ_1_) as a random effect, in order to account for differences between individual events in their tendency to predict gaze to the target talker. We further included an intercept for each participant (γ_2_) as a random effect, which allowed the model to account for differences between participants in their overall tendency to look at the target talker. We estimated the effect size of our models by looking at the variance explained by the fixed effects alone versus the complete model (including both fixed and random effects). This was done by calculating the pseudo-R-Squared value for Generalized Mixed-Effect model from the MuMIn package (https://cran.r-project.org/web/packages/MuMIn/MuMIn.pdf).

### Model 1: Effects of talker overlap and silence in turns and transitions

Prior to answering our research question, we first wanted to establish how the tendency to look at the active talker changed in events with overlap or silence during turns or transitions. Given that realistic conversation has many sources of potential variation to control for if it is to be used in an experiment, we wanted to know if conversation event type is a meaningful explanatory factor. This system of classification has the advantage that it can be automatically applied by examining the timeseries of voice activity, and is therefore more efficient than manual conversational analysis. Therefore, for our first model, the independent variable was event type and included five categorical levels (silence-transition, silence-turn, solo, overlap-transition, and overlap-turn). This was modelled as a fixed effect with the random effects described above (Equation 1). In addition, we included an independent slope parameter for each participant to account for differences between participants in how they behave for each event type as a random effect (γ_3_+Participant·Event Type). To evaluate the model, we used a likelihood ratio test to compare the full model in Equation 1 to a null model with the intercept fixed at 1.

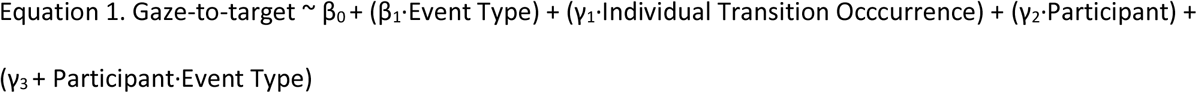

In our first model, we found that the odds of gazing towards the target talker during a solo were 2.21 (95% CI: [1.86, 2.63]). This model explained 38% of the variance, but only 0.5% was attributable to the effect of conversation event type. Including conversation event type did not significantly improve the model fit as compared to the null model (where the intercept was modelled as 1; likelihood ratio test: χ^2^_(4)_ = 8.1853, *n*.*s*.). This suggests that differences in conversation event type are not predictive of where listeners look when viewing a multitalker conversation. Nonetheless, the model allowed us to describe where a listener looked during each conversation event type as a probability with confidence intervals (Table 2). Furthermore, the model assessed how well each event type changed the odds of gaze-to-target, as compared to solos (Figure 3). Only during silence in a turn were the odds of gazing at the target talker significantly changed from the odds in a solo, where gaze-to-target was 1.77 times more likely (*p* < 0.001, 95% CI [1.29, 2.44]). Given that we found no evidence that any conversation event type decreased the odds of gaze-to-target, turns that included overlap and silence were not excluded from our subsequent analysis.

**Table 2.** Listeners’ gaze based on number of talkers (silence, solo, or overlap) and turn-taking stage (turn or transition).

| Event Type | Probability of gaze-to-target<br>[95% confidence interval] |
| --- | --- |
| Silence-Turn | 0.797 [0.854, 0.723] |
| Solo | 0.689 [0.725, 0.65] |
| Overlap-Turn | 0.654 [0.739, 0.558] |
| Silence-Transition | 0.749 [0.806, 0.681] |
| Overlap-Transition | 0.732 [0.769, 0.691] |

Legend: Values are calculated from multilevel logistic regression modelling and converted to probability from log odds. "Target" is defined as the turn-holding talker during turns, and as the finishing talker during transitions. Note that the confidence intervals were calculated with each event type, respectively, as the baseline intercept for the model.

**Figure 3.**
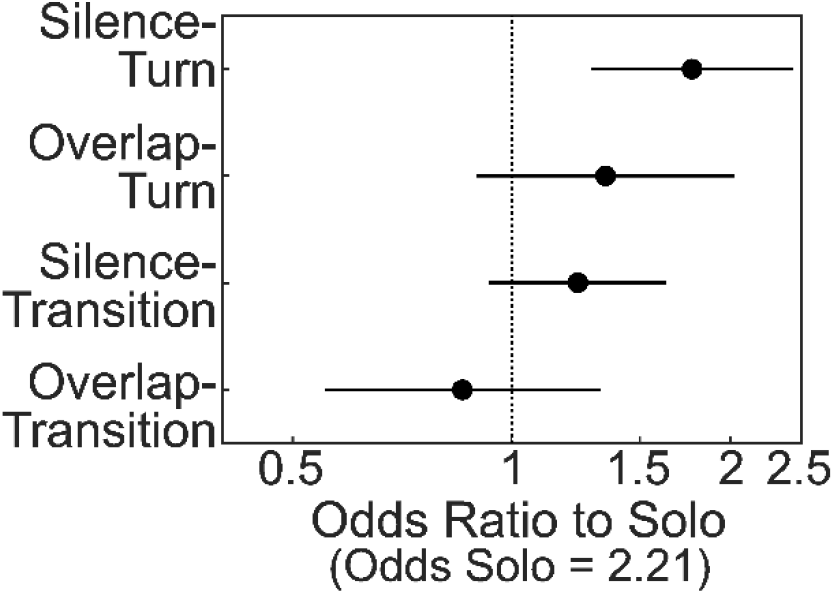
Change in odds of listener’s gaze-to-target from solo (1 active talker) to silence (0 active talkers) and to overlap (2 active talkers), in conversation turns and transitions. “Target” is defined as the turn-holding talker during turns, and as the finishing talker during transitions. Dotted line indicates odds that are equal to the odds in solo. Horizontal bars represent confidence intervals.

### Model 2: Effect of time on saccades during a talker transition

Our research question was with respect to when, and how much, hearing-impaired listeners look at the new talker during a conversational transition. To answer this, transition events identified from the conversation analysis were divided into time-bins of 100 ms duration. As such, each transition period had at least 5 time-bins, and up to 25 (time window start between 0.5-0.1 s before turn onset, and time window end between 0.4-2 s after). Thus, our independent variable consisted of time-bin, with 25 different levels, modelled as a fixed effect. We tested both a full model with the random effects outlined in the general modelling approach (Equation 2), and two reduced models with each of the random effect terms excluded, as well as a null model with the intercept fixed at 1. These were compared to the full model with likelihood ratio tests.

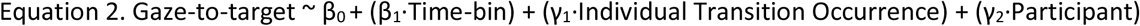

The odds of gaze-to-target in each time-bin are presented in Figure 4. The full model explained 38.7% of the variance, and the fixed effect of time alone explained 22.7% of the variance. The inclusion of time as a fixed effect improved the model fit compared to the null model (where the intercept was fixed at 1, likelihood ratio test: χ^2^_(27)_ = 21559, *p* < 0.001). This indicates that time over a talker transition is predictive of the listeners’ gaze towards the new talker. The presence of either random effect (random intercept for either participant, or for individual transition occurrence) improved the model fit (likelihood ratio tests, Eq. 2 vs. Eq. 2 excluding (γ_1_·Individual Transition Occurrence) term: χ^2^_(1)_ = 10943, *p* < 0.001; Eq. 2 vs. Eq 2 excluding (γ_2_·Participant) term: χ^2^ _(1)_= 1574.1, *p* < 0.001). Excluding a random intercept parameter for each participant only slightly decreased the variance explained by the model (from 38.7 to 36.9%, ΔAIC = 1572), whereas excluding that for the individual transition occurrences decreased the variance explained by more than half (from 38.7 to 23.1%, ΔAIC = 10941). This indicates that a substantial amount of the explanatory power of the model came from the fact that it accommodated differences in gaze behavior between individual transition events.

**Figure 4.**
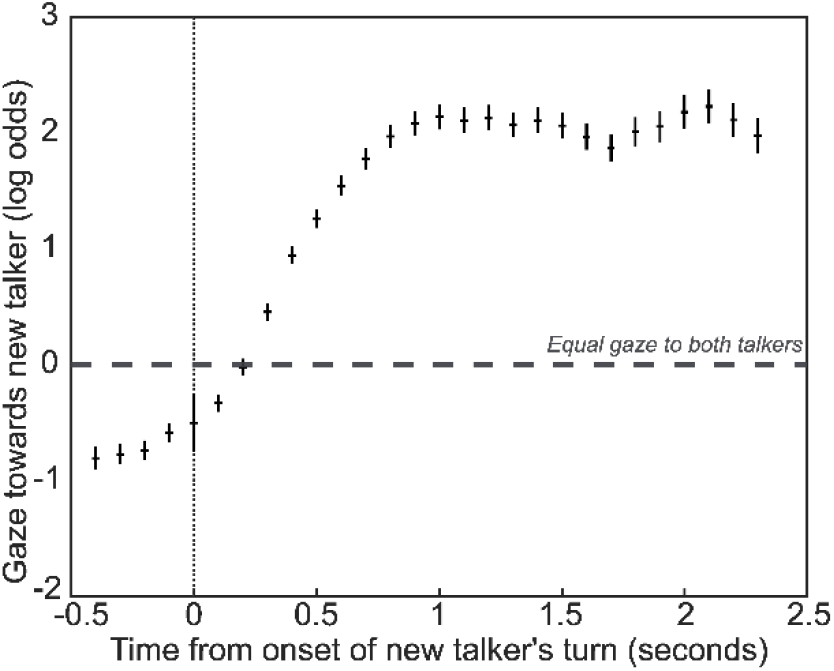
Odds of listener’s gaze to the new talker during a conversation transition. Solid vertical bars represent confidence intervals. Onset of new talker’s voice activity occurred at 0 seconds and was preceded by silence.

## Discussion

In the current investigation, we introduced a new paradigm for studying a listeners’ experience of a realistic multitalker conversation that is intended to capture the perception of turn-end cues. Using MLR modelling, we calculated the odds that a hearing-impaired listener will look towards a new talker as a function of time over a transition between talkers. Below we discuss these results, as well as the insights that we gathered from the process, in light of previous and future research.

### Timing of listener gaze during a transition between talkers

From the results of model 2, we observed that the likelihood of the listeners’ gaze towards the new talker peaked at 1 s after the new talker’s speech onset, with odds of 2.14 (Figure 4). These odds translate into a 68% chance that the listeners’ gaze was to the new talker. Table 3 summarizes previous research that has described the gaze trajectory of listeners with normal hearing, who similarly followed talker transitions in realistic conversation. The values reported in the current investigation are close to what has been observed in the normal-hearing population, with perhaps a longer timing to the peak of the gaze towards the new talker. We cannot draw conclusions from this comparison, however, due to substantial differences in methodology between the current and previous works. Notably, in our experiment, the conversation audio was accompanied by multitalker babble noise whereas in previous research the audio was always presented without noise.

**Table 3.** Six studies that report the gaze trajectory from listeners with normal hearing following talker transitions in audiovisual conversations.

|  | No.<br>Talkers | Peak |  | Conversation |
| --- | --- | --- | --- | --- |
|  |  | Probability of gaze to<br>new talker | Timing from onset of new<br>talker's speech |  |
| Dawson & Foulsham,<br>2022 | 3 | 55 | 450-500 ms | Unscripted |
| Foulsham &<br>Sanderson, 2013 | 4 | 60 | 829 ms | Unscripted |
| Hirvenkari et al.,<br>2013 | 2 | -- | *300 ms | Unscripted |
| Tice & Henetz, 2011 | 2 | <sup>v</sup>80 | <sup>v</sup>800 ms | Scripted,<br>Question-answer<br>exchanges |
| Casillas & Frank,<br>2012 | 2 | -- | <sup>β</sup>600 ms | Unscripted,<br>Question-answer<br>exchanges |
| Keitel et al., 2013 | 2 | -- | <sup>†</sup>750 ms | Scripted |

\*as reported in discussion, Hirvenkari et al. 2013.
<sup>v</sup> deduced from visual inspection of proportion of looks over time, Tice & Henetz (2011) Fig.1
<sup>β</sup> deduced from visual inspection of gaze direction over time, Casillas & Frank (2012) Fig.3
<sup>†</sup> deduced from visual inspection of frequency of anticipatory gaze shifts over time, Keitel et al. (2013) Fig. 4

The data presented in Figure 4 show a gentle increasing slope in the odds that the listener looked at the new talker over the silent period leading up to the onset of the new talker’s speech (from -0.4 to 0 s), which grew steeper in the first 300 ms after the talker’s speech began. Given that saccades towards a new talker require 200 ms to plan (Matin et al., 1993), changes in the odds of gaze to the new talker that occur within the first 200 ms of the new talker’s speech must have been initiated prior to the talker’s voice activity. As such, this early increasing slope may be interpreted as evidence of anticipatory gaze: Listeners were able to plan their saccade to the new talker prior to the onset of that talkers’ speech, despite that the log odds of gaze to the new talker were only positive at timepoints greater than 200 ms. Although previous research has varied in its definition of what constitutes anticipatory gaze, our interpretation is consistent with conclusions from some previous work in listeners with normal hearing (Casillas & Frank, 2012; Hidalgo et al., 2022; Holler & Kendrick, 2015; Keitel et al., 2013; Tice & Henetz, 2011). It is also mirrored in timeseries data from listeners with normal hearing that show an increasing slope within the first 200 ms after the onset of the new talker’s speech (Dawson & Foulsham, 2022; Foulsham & Sanderson, 2013; Hirvenkari et al., 2013).

### Listener’s gaze towards the turn-holder

Our results do not support that our classification of conversation event types according to the number of talkers (Figure 2) was predictive of where listeners look when viewing a multitalker conversation. Nonetheless, this analysis allowed us to quantify where a hearing-impaired listener looks during conversation (Table 2). Similar to previous research, we observed that listeners preferred to look at the active talker when only one talker was speaking, with odds of 2.21. This translates to 68.85% of the time. This value is in line with reports from previous research with listeners with normal hearing that used two-talker conversations without background noise, which include 63% (Casillas & Frank, 2012), 73.5 % (Hirvenkari et al., 2013), and 75% (Tice & Henetz, 2011) of the listeners’ gaze directed to the active talker. It is also close to the values that were reported for hearing-impaired listeners for conversations that occurred with a range of background noise levels (55-60%, based on visual inspection of (Lu et al., 2021) Fig. 6).

To what has previously been reported on a listener’s gaze during a multitalker conversation, in the current investigation we add a description of where listeners look during the more variable portions of realistic conversation, namely periods of overlap between the talkers and silence. There are no detailed reports on how these affect the listeners’ gaze, and this information is helpful for both the experimental design of research that uses realistic conversation stimuli, and for guiding applied research into how listener eye-gaze can be used as a control signal in hearing technology. We found that hearing-impaired listeners prefer to look at the turn-holding talker during turns, and the finishing talker during transitions, regardless of overlapping talkers or silence. There was an increase in the odds that the listener looked at the active talker during silence in a turn (Figure 3), which most likely reflects that gaze during a solo may have a gradual increase if the solo represents the beginning of a talker’s turn, whereas a silence is most likely occurring later in the turn after gaze towards the turn-holder has stabilized at its peak level. Pragmatically, these results suggest that in future research on eye-gaze in conversation, with stimuli that have a similar distribution of conversation event types as the current work (Table 1), potential effects from the number of active talkers can be disregarded.

### Mechanisms affecting listener gaze during a transition between talkers

Despite the congruence of the results from the current investigation with previous work on listeners with normal hearing, there is reason to believe that hearing-impaired listeners will show different eye-gaze patterns than this population when viewing a multitalker conversation. Specifically, listeners with normal hearing gaze more at the active talker than hearing-impaired listeners when the conversation occurs in high background noise (Lu et al., 2021). This difference between the populations may reflect their abilities to compensate for degraded audio – Listeners with normal hearing similarly show reduced gaze to the active talker when the audio is completely muted (Dawson & Foulsham, 2022; Foulsham & Sanderson, 2013; Hirvenkari et al., 2013; Tice & Henetz, 2011). At least three different mechanism could support this reduction in gaze in the hearing-impaired listeners. Specifically, it could be a reflection of: (1) the listener’s disengagement from the stimuli, (2) their difficulty determining which talker is active, possibly due to impaired turn-end perception, and/or (3) their slower reactions to changes in the active talker, due to the increased cognitive load of listening with hearing impairment. In theory, each of these mechanisms would also influence the s-shaped portion of the function shown from approximately -0.4 to 1 s in Figure 4, for example by flattening the peaks, shifting the x-axis crossing in time, or decreasing the slope. The sensitivity of our experiment paradigm to these changes remains a topic for future research.

### Assessment of the paradigm’s ecological validity

In the current investigation, our ultimate goal was to advance our understanding of how hearing-impaired listeners are challenged by communication in everyday conversations. This purpose aligns with the framework proposed for ecological validity in hearing research from the 6^th^ Eriksholm Workshop (purpose A, (Keidser et al., 2020). As a step towards this goal, we introduced a new paradigm that captures realism with both the choice of outcome measurement – eye-gaze, a largely automatic indicator of attention and cognition – and the choice of stimuli – life-sized, audiovisual recordings of an unscripted conversation. That being said, several factors that were essential to experiment control may have reduced the realism of the listener’s experience. These include the laboratory setting, the use of extensive measurement equipment, the trial-based presentation of the stimuli, and the requirement for the listener to complete a comprehension task.

A key difference between the stimuli used in the current experiment as compared to what a listener may experience in real-life listening was our use of recordings rather than live conversation with the listener’s participation. In real-life conversation, listeners use their eyes to communicate their attention and intentions to others (for a review, see Degutyte & Astell, 2021). Real talkers, rather than recorded ones, would better invite the potential for interaction, and may therefore elicit different eye-gaze behavior than that observed here. This potential for interaction also changes the cognitive processes involved in listening, as it invokes more complex neural systems than those involved in passive observation alone (for a review see Carlile & Keidser, 2020). On the other hand, the use of recordings in the current investigation allowed us to assess how reliable eye-movements were between participants and between individual transition events. Our results suggest that listeners behave more similarly when viewing the same conversation stimulus than they do from one stimulus to the next. As such, one may expect worse accuracy at predicting gaze when stimuli are not repeated across test participants, such as in an experimental setup where the conversation is live. Thus, for the purpose of experimental control and consequent experimental sensitivity, using recorded stimuli may have advantages.

### Future Outlook

We intend for the current paradigm to be used as a starting point for future work on hearing in realistic conversations. At the forefront of today’s research, scientists are designing experimental setups that fuse information from numerous sensors to generate rich multimodal datasets, capable of capturing multiple behaviors from realistic communication situations (e.g. Aliakabary et al., 2022; Donley et al., 2021; Fredriksson & Wallin, 2020; Hadley et al., 2019; Lu et al., 2021). These complex data will benefit from concrete *a priori* hypotheses and analysis plans in order to extract meaningful signals, for which we hope that the proposed paradigm offers a justified starting place. Furthermore, recent advances in portable eyetracking, such as methods for smartphone camera-based algorithms (e.g. Valliappan et al., 2020) and in-ear electrooculophraghy (e.g. Skoglund et al., 2022), open possibilities for collecting data outside of the traditional laboratory, for example in an audiologists’ clinic or in the hearing-aid users’ real life. These possibilities mean that future research can cover a wider population and/or situations that are increasingly realistic. Given the growing interest in the behavior of hearing-impaired listeners in realistic settings, we hope that this work will expedite progress towards more ecologically-valid research outcomes in the field of hearing science.

## Acknowledgements

This research was supported in part by the Swedish Research Council (Vetenskapsrådet, VR 2017-06092 418 Mekanismer och behandling vidåldersrelaterad hörselnedsättning). All authors are employed by Oticon A/S (Smørum, Denmark). The authors would like to acknowledge Martin Andersen and Mike Lind Rank (T&W Engineering A/S, Allerød, Denmark) for their contributions to the experiment from which we resourced the data of the current investigation (Skoglund et al., 2022).

